# An empirical test of the temperature dependence of carrying capacity

**DOI:** 10.1101/210690

**Authors:** Joey R. Bernhardt, Jennifer M. Sunday, Mary I. O’Connor

## Abstract

Predicting population persistence and dynamics in the context of global change is a major challenge for ecology. A widely held prediction is that population abundance at carrying capacity decreases with warming, assuming no change in resource supply, due to increased individual resource demands associated with higher metabolic rates. However, this prediction, which is based on metabolic scaling theory (MST), has not been tested empirically. Here we experimentally tested whether effects of temperature on short-term metabolic performance (rates of photosynthesis and respiration) translate directly to effects of temperature on population rates in a phytoplankton species. We found that effects of temperature on organismal metabolic rates matched theoretical predictions, and that the temperature dependence of individual metabolic performance translated to population abundance. Population abundance at carrying capacity, *K*, decreased with temperature less than expected based on the temperature dependence of photosynthesis. Concurrent with declines in abundance, we observed a linear decline in cell size of approximately 2.3% °C^−1^, which is consistent with broadly observed patterns in unicellular organisms, known as the temperature-size rule. When theoretical predictions include higher densities allowed by shifts toward smaller individual size, observed declines in *K* were quantitatively consistent with theoretical predictions. Our results indicate that outcomes of population dynamics across a range of temperatures reflect organismal responses to temperature via metabolic scaling, providing a general basis for forecasting population responses to global change.

## Introduction

Understanding population persistence and dynamics in a changing environment is a major challenge in ecology. Population dynamics reflect individual organisms’ performance, which can change with temperature and affect demographic vital rates and ultimately population persistence (Fridley 2017). Despite substantial theoretical and empirical evidence linking temperature to one key demographic parameter - the intrinsic growth rate *r* - there has been little attention given to how changing temperature affects another central population parameter: density at steady state, or carrying capacity, *K* (Savage et al 2004, O’Connor et al. 2011, Gilbert et al. 2014). Carrying capacity of resource populations is central to understanding the structure of ecosystems and their stability (Rosenzweig 1971). In the absence of empirical evidence on the relationship between temperature and *K*, some models have assumed that *K* declines with increasing temperature proportionally to temperature-induced increases in per capita resource use (Allen et al. 2007, O’Connor et al. 2011, Gilbert et al. 2014). Yet different assumptions about how *K* changes with temperature have led to different predictions about the ecological outcomes of warming (e.g. Osmond et al. 2017 vs. Sentis et al. 2017). To date, we still lack an empirical test of whether population carrying capacity declines with temperature, and if so, if that decline should be predicted by temperature-driven change in per capita metabolic rate.

Carrying capacity, *K*, is the non-zero population abundance at which population growth is equal to zero. *K* is not merely a theoretical endpoint of the logistic model at stable equilibrium (Equation 1); it is one of two central parameters that describe temporal patterns in population abundance in many dynamic models. Although simple, the logistic growth model effectively describes population growth in microbial populations in simple environments, and underlies more complex models. In the logistic growth model,

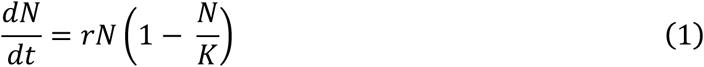

*N* is the size of the population, and carrying capacity, *K*, is the value of *N*>0 that makes *dN/dt* = 0 (Verhulst, 1838, Gotelli 1995).

A population’s carrying capacity, *K*, is the outcome of density-dependent population growth (Gause 1932, Gotelli 1995). The strength of density-dependence, or the effect of intraspecific competition in limiting population growth at high densities, determines population abundance when it is near *K*, and reflects the impact of density on per capita rates of resource use, birth and death. These rates vary with the temperature dependence of metabolic rate in similar ways across diverse taxa (Gillooly et al. 2001, Dell et al. 2011, Pawar et al. 2016). Metabolic scaling theory (MST) has postulated that this variation is due to highly conserved, temperature-dependent rates of aerobic respiration and oxygenic photosynthesis that underlie resource use, growth and survival rates, lending some predictability to temperature effects on demographic processes (Brown et al 2004, Savage et al 2004, O’Connor et al. 2011). Empirical evidence for the temperature dependence of per capita rates and population growth rates supports MST predictions (Eppley 1972, Ernest et al. 2003, Savage et al. 2004, López-Urrutia et al. 2006, López-Urrutia 2008). Still, there is no evidence for how temperature dependence of per capita performance affects density when density dependence is strong, near carrying capacity. Further, no experiments have tested whether the macro-ecological relationships between the temperature dependence of per capita photosynthetic or respiration rates and abundance at carrying capacity hold at the scale of a single population under controlled conditions. This is a major gap between the macro-ecology of metabolic scaling theory and the empirical patterns observed at the scale of warming experiments and study sites.

To solve this problem, we consider a general model for how metabolic thermal constraints translate to density dependence and abundance at carrying capacity (Savage et al. 2004). We estimate how per capita performance changes with temperature and whether that change directly predicts carrying capacity across a thermal gradient. We first test the hypothesis that the temperature dependence of per capita metabolic rate (oxygen flux) accurately predicts the decline in abundance with temperature. We then consider concurrent temperature-related shifts in phenotype, in particular, changes in body size consistent with the widely-observed temperature size-rule (Atkinson et al. 2003). We express these hypotheses and findings mathematically and integrate them into the Savage et al. (2004) model of how temperature dependence of metabolism scales to population vital rates.

## Methods

To address these questions, we express our empirically testable hypotheses in terms of the Savage et al. (2004) model, which links the temperature dependence of metabolism to classic population growth model parameters (*r, K*). The Savage et al. (2004) model is centered on the allocation of energetic resources to processes that affect demography: survival, growth, and reproduction. We highlight the allocation assumptions here because they underlie our present approach. We describe how this theory can be tested empirically with independent measures of metabolism and population dynamics and how this theory might be combined with the temperature-size rule (Atkinson et al. 2003, Forster et al. 2012) to predict changes in *K.* We then outline two resulting hypotheses that we test empirically with phytoplankton populations.

### Deriving experimentally testable hypotheses

Populations can be maintained at steady state when individuals are reproducing and births balance deaths at the population level, maintaining constant density (births = deaths ≠ 0). Energy (*E*) consumed by individuals must be allocated to producing new individuals and to maintenance. Following Savage et al. (2004), if *E*(*M, T*) is the mass-(*M*) and temperature-(*T*) dependent per capita energy required to produce a new individual, then it takes *N*(*M, T, t*)*E*(*M, T*) amount of energy to replace the entire population, where *N*(*M, T, t*) is the number of individuals at time *t*. At carrying capacity, on average *N* deaths will occur over a time period equal to the average lifespan, *S*(*M, T*), and all individuals will be replaced over a time period equal to the average lifespan, so the energy needed to keep the population size at steady state is *N*(*M, T, t*)*E*(*M, T*)*/ S*(*M, T*). If we assume that energy required to produce a new individual (*E*(*M*)) is linearly related to its mass, and independent of temperature (i.e. temperature may affect the rate of ontogenetic growth, but not the energy required to produce a new individual), then the total metabolic rate of the population (B_*pop*_) at steady state (*N*=*K*) is

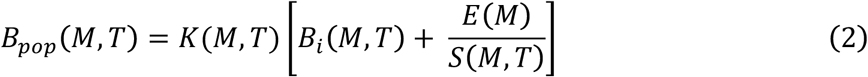

where, after expansion, the first term is the energy required for maintenance and the second term, *K*(*M, T*)*E*(*M*)*/S*(*M, T*), is the energy required for replacement (i.e. production of one new individual per individual) and *B*_*i*_(*M*, *T*) is individual metabolic rate. Assuming that total resource use by the population equals total metabolic rate of the population, (B_*pop*_), then carrying capacity is achieved when the rate of resource supply, *P*, in the environment equals the rate of resource use by the population (B_*pop*_). If *S*(*M*, *T*) = *S*_*0*_/*B*_*i*_(*M,T*) (i.e., lifespan scales inversely with per capita metabolic rate, Gillooly et al. 2001) and we assume that *E*(*M*)/*S*(*M, T*) = (*E*_*0*_/*S*_*0*_)*B*_*i*_(*M, T*) then,

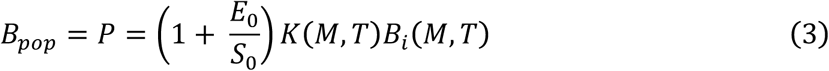

where *E*_*0*_ and *S*_*0*_ are mass- and temperature-independent normalization constants, *K*(*M, T*) is population abundance at steady state, and *B*_*i*_(*M, T*) is individual metabolic rate (Equation 11 in Savage et al. 2004).

Equation 3 can be rearranged to show that, when resource supply (*P*) is constant and independent of temperature, the temperature and mass dependences of *K* and *B*_*i*_ at steady-state must be inversely proportional, leading to the prediction that carrying capacity declines as individual metabolic rate increases with temperature (Equation 4). When individual metabolic rate, *B_i_,* scales with mass and temperature as *B*_*i*_ = *M*^*3/4*^*e*^−*Ea/kT*^ (West et al. 1997, Gillooly et al. 2001), and if we assume for now that mass is independent of temperature, then *K* is predicted to decrease with increasing body size and decrease with increasing temperature (Savage et al. 2004; Figure 2):

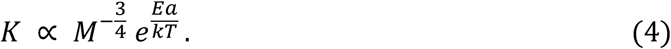

Therefore, a reasonable prediction for carrying capacity in warming environments is that it should decline with the slope of the inverse temperature dependence of per capita metabolic rate (Savage et al. 2004, Vasseur and McCann 2005, O’Connor et al. 2011, Gilbert et al. 2014). This prediction assumes, however, that body size does not depend on temperature, so *K* scales simply as a function of the activation energy of metabolism (i.e. *K* is proportional to *e^Ea/kT^*). This simplifying assumption contradicts evidence for widely observed declines in body size associated with warming (Atkinson et al. 2003). Changing body sizes with warming could alter predictions of population density (*K*) as a function of temperature. Incorporating temperature-dependent body size into expectations of population abundance, as we do next, may more closely link theory for temperature effects with observed changes in experiments and nature.

Following Osmond et al. 2017, we model the hypothesis that a temperature-induced decline in *K* is modified by concomitant changes in body size by modifying Equation 4 to allow individual body mass to depend on temperature. We assume that *M*(*T*) declines linearly with temperature (consistent with the temperature-size rule in ectotherms, TSR),

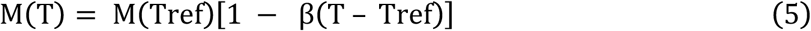

where β is the fraction that mass is reduced as temperature is increased by one degree, and *Tref* is a reference temperature (here 5°C). This linear approximation of the TSR is appropriate for unicellular organisms such as phytoplankton (Atkinson et al. 2003, DeLong 2012, Forster et al. 2012). If body mass decreases with temperature, then the negative temperature dependence of *K* should be reduced relative to the case where body mass is temperature-invariant, because warmer conditions should support relatively more individuals of smaller size, thus reducing the negative temperature dependence of *K*. Alternatively, if body size does not change with temperature (β= 0), then the temperature dependence of *K* should be inversely proportional to the activation energy of metabolism, as shown above (Equation 4).

Here, in a closed phytoplankton mesocosm system with a fixed, finite nutrient supply, we experimentally tested the following two hypotheses:

*Hypothesis 1:* Carrying capacity declines with increasing temperature proportionally to the activation energy of per capita metabolic rate over a range of non-stressful temperatures. For primary producers, such as phytoplankton, the temperature dependence of *K* varies as a function of the activation energy of photosynthesis (*E*_*a*_ ≈ 0.32 eV) (Allen et al. 2005, Dewar 1999, Lopez-Urrutia 2006, 2008) and body mass at a reference temperature (Equation 4). At higher temperatures, per capita metabolic demand increases following a Boltzmann-Arrhenius relationship and causes carrying capacity, *K*, to decline. This relationship requires that total population level resource use does not increase such that the effect of temperature has the potential to translate to per capita resource limitation (Figure 1).

**Figure 1.**
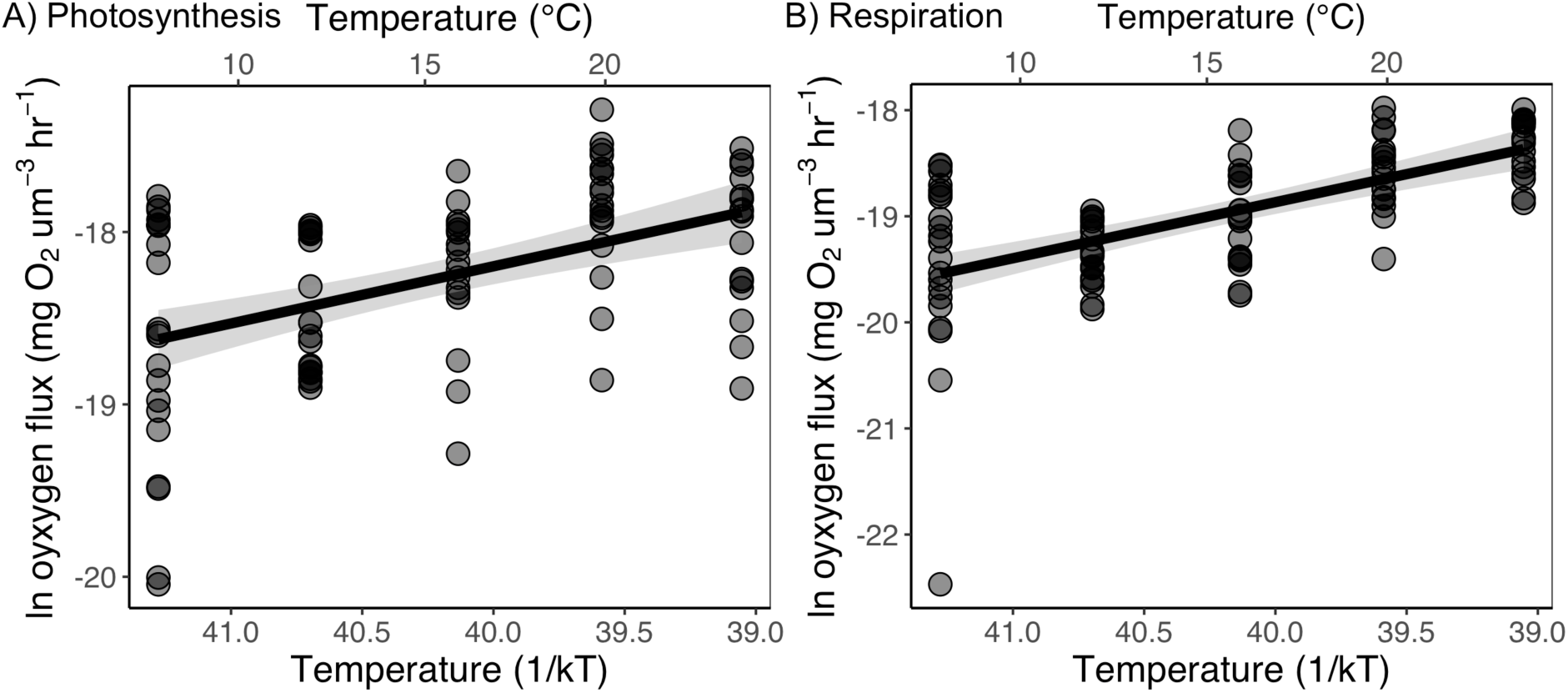
Mass-normalized metabolic rates of *T. tetrahele* increase with temperature. Estimated activation energies are for gross photosynthesis *Ea* = 0.33 (95% CI: 0.28, 0.37) (A), and for respiration, *Ea* = 0.53 (95% CI: 0.39, 0.67) (B). Points are shown at medium opacity to indicate overlap.

*Hypothesis 2:* Body mass decreases with temperature consistent following the temperature-size rule (Atkinson et al. 2003), reducing the effect of temperature on density at carrying capacity (Equation 5).

### Experimental Methods

*Tetraselmis tetrahele* is a globally distributed coastal marine phytoplankton species. The cultured strain used here was obtained from the Canadian Centre for the Culture of Microorganisms (UW414), and was originally isolated off the coast of Vancouver Island, British Columbia, Canada. *T. tetrahele* were maintained in laboratory culture in ESAW medium (Enriched Seawater, Artificial Water, Harrison et al. 1980) in 30 mL glass test tubes at 16°C for one year on a 16:8 light:dark cycle under nutrient and light saturated conditions before the start of the experiments. *T. tetrahele* is a flagellated chlorophyte that is fast-growing, eurythermal, highly motile, with a generation time of less than a day (Pena and Villegas 2005), making it a suitable species for mesocosm studies and tests of metabolic scaling theory.

### Estimating the activation energy of photosynthesis

We determined the activation energy of photosynthesis over a temperature range from 8°C - 24°C by measuring oxygen evolution in the light and oxygen consumption in the dark using a 24-channel optical fluorescence oxygen system (Sensor Dish Reader SDR2, PreSens), equipped with a 24-chamber 200 uL glass microplate (Loligo Systems Aps, Tjele, Denmark). The reader was placed in a temperature-controlled incubator (Panasonic M1R-154) with light at 80 umol/m^2^/s over the course of the experiments. Prior to measurements of metabolic rates, 200 uL of well-mixed *T. tetrahele* cultures were transferred from 30 mL test tubes to each well on the microplate. Wells were sealed with transparent PCR film (Thermo Scientific, Waltham, MA, USA), and measurements of oxygen concentrations were taken every 15 seconds over three hour periods, first in darkness and next in light, using the SDR v4.0 software (PreSens, Germany). Prior to oxygen flux measurements, sensor spots were calibrated with air-saturated water and water containing 2% sodium sulfite at each experimental temperature. Phytoplankton cells were acclimated to the assay temperature for an hour in the dark prior to measurements. Six blank wells containing ESAW medium were run at the same time as the phytoplankton, and the average rate of oxygen flux in these wells was subtracted from the experimental wells to account for background microbial respiration. Gross photosynthesis (GP) was estimated as GP = net photosynthesis + respiration at each temperature. We assumed that net photosynthesis is directly proportional to oxygen production in the light. We estimated per capita mass-normalized metabolic rates (*B*_*i*_) by dividing the total oxygen fluxes by the total population biovolume (mean cell volume * cell density) from the source cultures measured using a FlowCAM (FlowCAM VS Series, Fluid Imaging Technologies) at a flow rate of 0.3 ml/min immediately before the respirometry experiments. The activation energies (*E*_*a*_, Equation 4) of gross photosynthesis and respiration were estimated from relationships between log transformed mass-normalized oxygen flux rates and temperature (1/kT) using OLS linear regression.

### Estimating the temperature dependence of carrying capacity

We initiated five replicate experimental populations of *T. tetrahele* in 30 mL glass test tubes containing 25 mL of 10uM nitrate ESAW medium at a density of 2000 cells/mL at 5°C, 8°C, 16°C, 25°C, 32°C, and 38°C. Nitrate concentrations in the medium were reduced (to 10uM) relative to other nutrients to ensure a controlled limiting nutrient at carrying capacity. Mesocosms were held at constant temperature and light conditions (16:8h light:dark cycle; 60 umol/m^2^/s) until they reached steady state at all temperatures. Cell densities (cells/mL) and biovolumes (um^3^/mL) were measured from 250 uL samples every four days at the same time of day using the FlowCAM (flow rate 0.3 ml/min) for 43 days, until populations at all temperatures had reached carrying capacity (i.e. steady state).

To compare empirical observations with the predictions derived from Savage et al.’s framework (Equation 4), we estimated *K* in terms of number of individuals (individual cells/mL; units consistent with the units of the Savage model) and in terms of population biomass (approximated as total biovolume, um^3^/mL). Population biomass is not explicitly modeled in Savage et al. (2004), and we add this measure here because it integrates population abundance and body size. We used a differential equation solver (*fitOdeModel* function with the ‘PORT’ algorithm in the *simecol* package in R) to fit a logistic growth model (Equation 1) to our time series population abundance data and estimated the parameters *r* and *K*. We set the initial phytoplankton abundance to our experimental starting conditions and examined model fits graphically by comparing simulated data using estimated parameters with the observed time series of population size. To test our prediction of the response of *K* across temperatures using the activation energy of photosynthesis in the increasing part of the thermal performance curve only (thus excluding thermally stressful conditions past the thermal optimum), we chose to restrict our analysis to temperatures up to and not exceeding the thermal optimum for intrinsic growth rates in *T. tetrahele* (Pawar et al. 2016).

To assess how population-level resource use varies with temperature, we created controlled resource conditions in the mesocosms. We reduced the concentration of nitrate in the medium by 55-fold relative to complete ESAW medium to ensure that nitrate concentrations were limiting population densities at carrying capacity. We confirmed that populations were nitrate-limited by comparing abundances in populations at steady state when grown at higher nitrate levels in pilot studies prior to the experiment. We ensured that light was not limiting by observing no increase in abundance at higher light levels. To assess how population-level nitrate use at carrying capacity changes with temperature, we measured the nitrate remaining in the mesocosm water columns at steady state.

At the end of the experiment, we measured chlorophyll-a concentration on a Turner Designs Trilogy fluorometer after filtration of 2mL samples of the experimental volume containing well-mixed phytoplankton onto GF/F filters, freezing the filters at -20°C and later extracting with 90% acetone. We assayed nitrate concentrations spectrophotometrically from the filtrate using a cadmium reduction method (Strickland and Parsons 1968; LaMotte Nitrate Nitrogen Test Kit) with a Turner Designs Trilogy fluorometer (Nitrate/Nitrite Module (P/N: 7200-074)). We conducted all statistical analyses in R (version 3.4.1) (R Core Team 2017).

## Results

*Hypothesis 1:* The activation energy of mass-normalized photosynthesis in *T. tetrahele* was 0.33 eV (95% CI: 0.20, 0.46) (Figure 1). Including the 32°C and 38°C populations in the activation energy estimation introduced a non-linear decline in ln *K*, consistent with the thermal optimum for *T. tetrahele* (approximately 28°C; *personal observations*), and therefore these populations were not included in the linear fit. At steady state, the natural log of population abundances (carrying capacity, ln *K*) decreased with increasing temperature at a rate of -0.18 eV (95% CI: - 0.24, -0.12) (Figure 2). This corresponds to a temperature dependence that is less than that expected under Hypothesis 1, which predicts a slope that is inversely proportional to the activation energy of photosynthesis (-0.33 eV, i.e. -0.18 eV > -0.33 eV). Population-level nitrate use at steady state did not change with temperature (slope = 0.055 eV, 95% CI: -0.29, 0.19) (Figure 3A).

**Figure 2.**
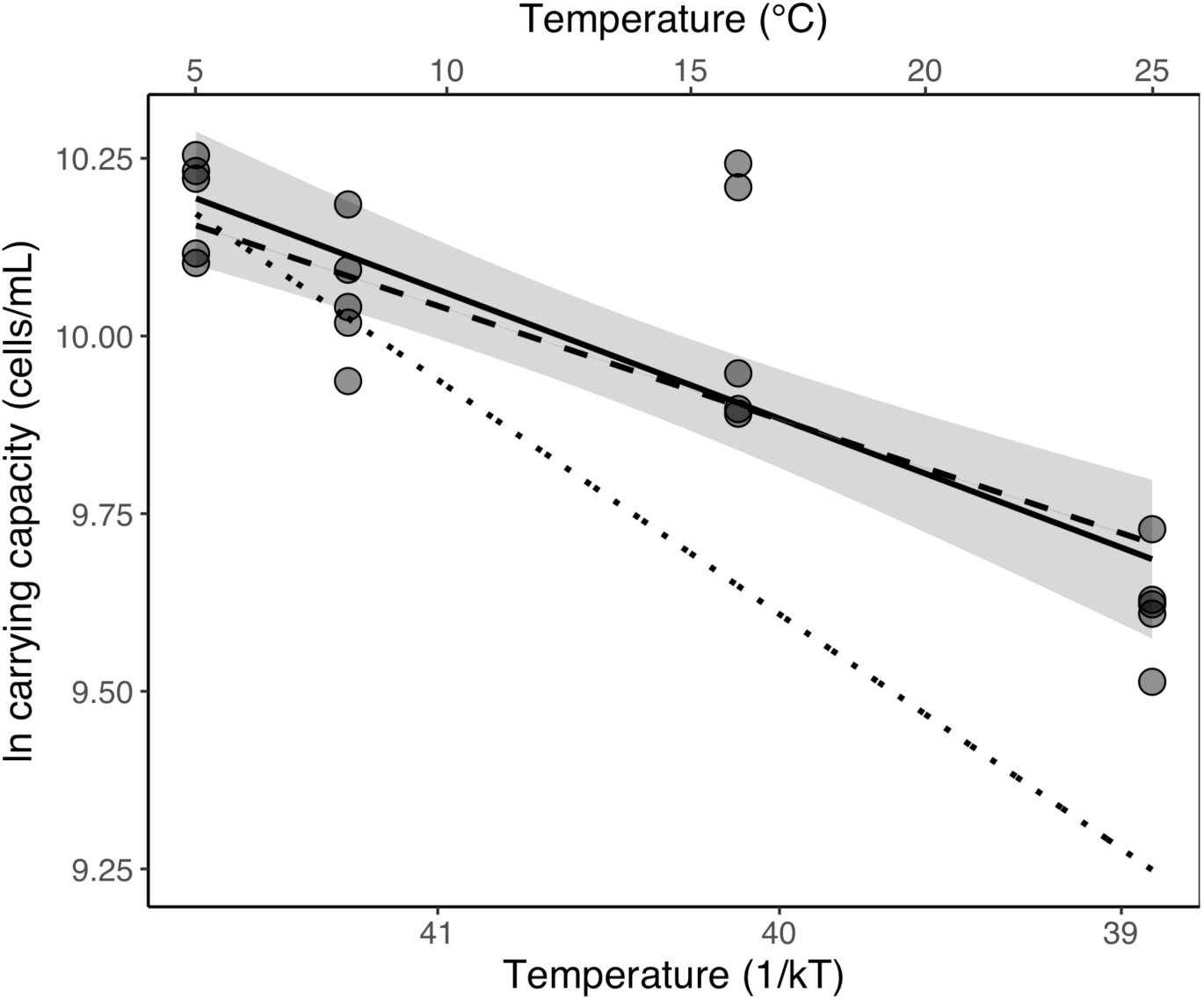
Carrying capacity decreases with temperature, with a slope of -0.18 eV (95% CI: -0.24, -0.12). Dotted line corresponds to the predictions of Savage et al. (2004), with a temperature-independent body mass. Thick dashed line corresponds to predicted carrying capacity, with a temperature-dependent body mass. Solid line corresponds to the linear model fit to data.

**Figure 3.**
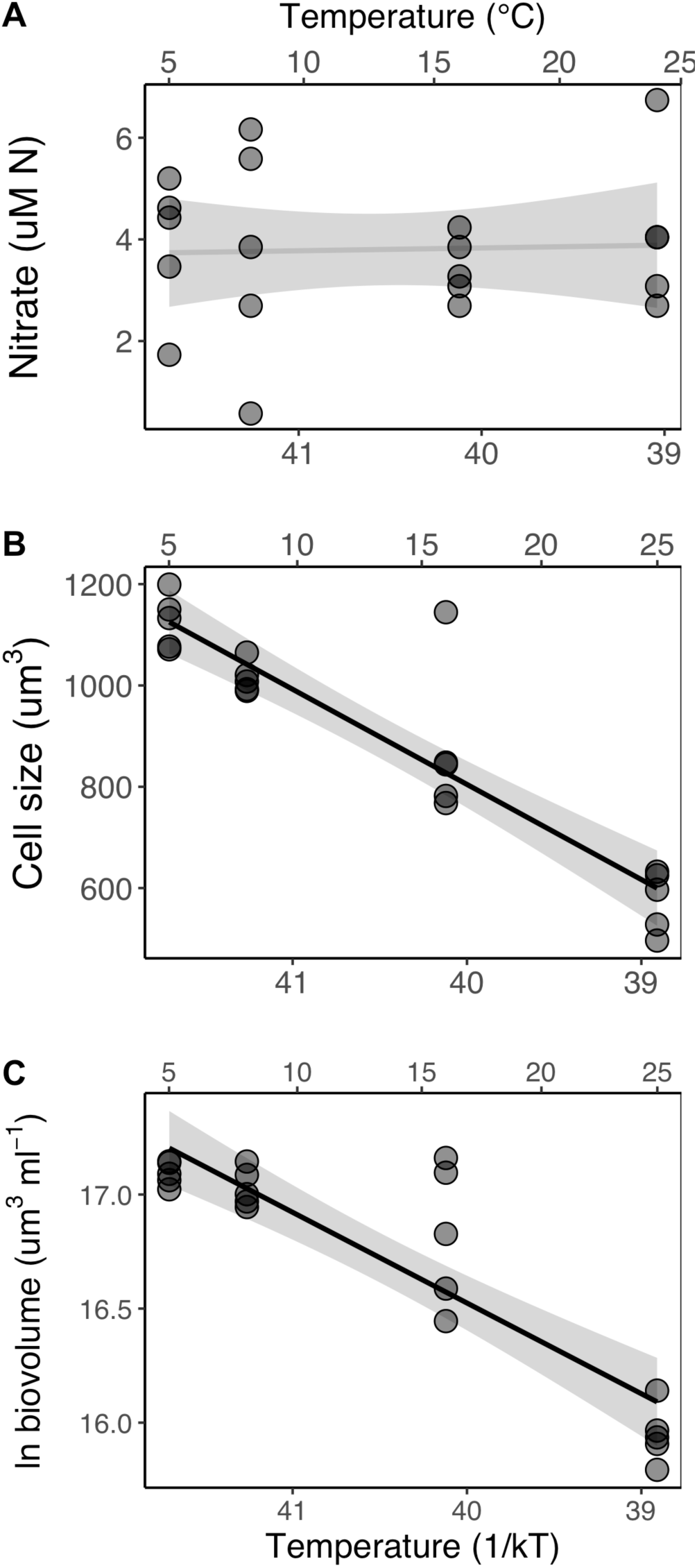
Water column total nitrate (A), average cell size (B) and total population biovolume (C) at steady state (after 43 days of growth).

*Hypothesis 2:* Cell size decreased as temperature increased (slope = -17.34 um^3^/°C, 95% CI - 20.71, -13.99) (Figure 3B), which corresponds to ~2.3% decrease in cell size per degree increase in temperature. When this observed decline in body size was included in the theoretical prediction for *K* (Equation 5), the predicted slope was -0.16 eV, which was statistically indistinguishable from the empirical estimates of *K* (-0.18 eV, 95% CI: -0.24, -0.12; Figure 2). Consistent with these patterns of declines in final abundance and size with temperature, carrying capacity estimated as population biomass, which combines estimates of cell size and cell number, decreased with temperature with a slope of -0.33 eV (95% CI: -0.37, -0.28) over the range of temperatures from 5°C - 25°C (Figure 3C). Population-level concentrations of chlorophyll-a also decreased with increasing temperature (slope = -0.67 eV, 95% CI -0.81, - 0.53).

## Discussion

Consistent with metabolic scaling theory and macro-ecological synthesis (Savage et al. 2004), we found that at the scale of single populations of *Tetraselmis tetrahele*, carrying capacity varies with the temperature dependence of photosynthesis and the temperature dependence of body size. We observed a linear decline in cell size of approximately 2.3%°C^−1^, which is consistent with broadly observed patterns in unicellular organisms (Forster et al. 2012). While *K* declined with warming, the concomitant reduction in body size meant that *K* did not decline by nearly as much as would have been predicted by MST when assuming a temperature-invariant body size. Body size shifts consistent with the TSR effectively compensated for declines in density that were expected based on metabolic demand.

To our knowledge, this is the most direct evidence to date for the proposed links between population abundance at carrying capacity, body size and empirically derived estimates of metabolic rate activation energies, thus providing a robust test of metabolic scaling theory at the population level. The estimated activation energy of photosynthesis in this study, 0.33 eV, is consistent with previously published estimates of the activation energy of photosynthesis in phytoplankton (López-Urrutia et al. 2006, Regaudie-de-Gioux and Duarte 2012, Padfield et al. 2016). Carrying capacity results here are qualitatively consistent with other studies that have found similar negative temperature dependence of carrying capacity (Alto and Juliano 2001, West and Post 2016), although the link between per capita metabolic rate and density was previously assumed rather than measured.

Carrying capacity declined more rapidly than expected at temperatures exceeding the thermal optimum in this species (i.e. the 32°C and 38°C populations). This decline is consistent with dominance of physiological stress responses to temperature as it rises past the thermal optimum. When abundance declines due to physiological stress associated with warming, ecological opportunities for invasion, species turnover, or adaptation are expected to occur and shift community function (Sorte et al. 2010).

Fundamental constraints on metabolism are reflected in the scaling of population density with body size and temperature (Enquist et al. 1998, Savage et al. 2004). The relationship between carrying capacity and temperature depends on how population-level resource use changes with temperature. Here we observed that population-level resource use (measured as nitrate remaining in the mesocosms at steady state) was the same across all temperatures, despite higher standing biomass and larger cell sizes in the cold. This suggests that under cold conditions, *T. Tetrahele* is more efficient at converting the limiting nutrient into biomass. In marine phytoplankton, nutrient uptake and conversion efficiency are strongly dependent on cell size. Maximum nutrient uptake rates increase isometrically with cell size (Marañón et al. 2013), while minimum nitrogen requirements scale with negative allometry (i.e. scale with a slope of ~0.87), meaning that larger cells are more mass-efficient at converting nutrients to biomass (Marañón et al. 2013). While nutrient uptake, use and efficiency are often dependent on cell size, limited empirical evidence suggests that per capita nitrate uptake rates are independent of temperature in at least one species of marine phytoplankton, *T. pseudonana* (Baker et al. 2016). Taken together, our observations are consistent with these patterns of increased nutrient use efficiency at larger cell sizes and suggest that populations of larger cells may be able to maintain higher population biomass under conditions of nutrient limitation.

Carrying capacity of primary producers is a central parameter used in consumer-resource models. To date, most studies of consumer-resource dynamics have assumed a negative temperature dependence of resource carrying capacity (O’Connor et al. 2011, Rall et al. 2012, Gilbert et al. 2014). Where nutrient supply is consistent across temperatures and the population is well mixed or highly mobile, ensuring equal access to resources, our observations of decreasing abundance with warming suggest that this is a valid qualitative assumption. However, more empirical work would help to justify its general use, given that at least one other study (DeLong and Hanson 2011) found a contrasting result. Our results further show that including the scaling of body size with temperature – even using the average response of approximately 2% decline per degree C– may generally improve accuracy.

Relating our findings to the patterns seen in other laboratory and field observations investigating population abundance as a function of temperature is not straightforward because most experiments have not explicitly fixed resource supply across temperature treatments. As a result, abundance changes with temperature are somewhat difficult to interpret in terms of energy-based predictions, and observed relationships have been increasing, decreasing, or unimodal in previous studies in which resource supply was not controlled (Fox and Morin 2001, Jiang and Morin 2004, Isaac et al. 2011).

Here we showed that carrying capacity and body size of photosynthetic autotrophs decline with increasing temperature, demonstrating a clear link between the kinetics of organismal metabolic rate and a key demographic parameter, *K*. We extended predictions of MST to include predictions that account for concomitant changes in body size with temperature – the temperature-size rule. We found that the temperature dependence of population abundance at steady state can be predicted based on changes in individual resource demand and body size with warming, thus demonstrating a metabolic basis of population dynamics. This work reinforces the framework of metabolic scaling of temperature dependence from subcellular processes to ecosystem processes, via population dynamics, for understanding cross-scale consequences of temperature in ecological systems.

## Acknowledgements

We thank J. Yangel and W. Ou for help with lab experiments. We’re grateful to M. Tseng, N. Knight, M. Siegle, N. Caulk and other members of the O’Connor Lab for many useful discussions about carrying capacity; and M. Osmond for feedback on the manuscript. Funding was provided by a Vanier Canada Graduate Scholarship (J.R.B), Natural Sciences and Engineering Research Council (M.M.O) and the Biodiversity Research Centre (J.M.S).

## Data accessibility

All data will be made available on Dryad should the manuscript be accepted.

## Author contributions

JRB conceived and designed the study, along with help from MMO and JMS. JRB carried out the experiments, analyzed the data and wrote the manuscript. All authors contributed to writing the manuscript.

